# Construction of a solid Cox model for AML patients based on multiomics bioinformatic analysis

**DOI:** 10.1101/2021.09.15.460430

**Authors:** Fu Li, Jiao Cai, Jia Liu, Shi-cang Yu, Xi Zhang, Yi Su, Lei Gao

## Abstract

Acute myeloid leukemia (AML) is a highly heterogeneous hematological malignancy. The bone marrow (BM) microenvironment in AML plays an important role in leukemogenesis, drug resistance and leukemia relapse. In this study, we aimed to identify reliable immune-related biomarkers for AML prognosis by multiomics analysis. We obtained expression profiles from The Cancer Genome Atlas (TCGA) database and constructed a LASSO-Cox regression model to predict the prognosis of AML using multiomics bioinformatic analysis data. This was followed by independent validation of the model in the GSE106291 (n=251), GSE12417 (n=163) and GSE37642 (n=137) datasets and mutated genes in clinical samples for predicting overall survival (OS). Molecular docking was performed to predict the most optimal ligands to these hub genes. The single-cell RNA sequence dataset GSE116256 was used to clarify the expression of the hub genes in different immune cell types. According to their significant differences in immune gene signatures and survival trends, we concluded that the immune infiltration-lacking subtype (IL type) is associated with better prognosis than the immune infiltration-rich subtype (IR type). Using the LASSO model, we built a classifier based on 5 hub genes to predict the prognosis of AML (risk score = −0.086×ADAMTS3 + 0.180×CD52 + 0.472×CLCN5 - 0.356×HAL + 0.368×ICAM3). In summary, we constructed a prognostic model of AML using integrated multiomics bioinformatic analysis that could serve as a therapeutic classifier.

## Introduction

Acute myeloid leukemia (AML) is a highly heterogeneous group of hematological malignancies that are characterized by various cytogenetic and molecular heterogeneities (Dohner, Weisdorf, & Bloomfield, 2015; Short, Rytting, & Cortes, 2018). Although substantial progress has been achieved with combinatorial therapies including radiation, chemotherapy, immunotherapy and/or targeted therapy, the cure rate of patients remains only 35%-40% in younger patients (age< 60 years) and 5%-15% in older patients (age< 60 years)(Döhner et al., 2010). Relapse and refractory disease continue to be major obstacles in the treatment of AML, with 29% or fewer patients living beyond 5 years.

Several studies have shown that changes in the bone marrow (BM) microenvironment in AML largely promote distinct biological processes in leukemogenesis, drug resistance and leukemia relapse(Ghobrial, Detappe, Anderson, & Steensma, 2018). Thus, insights into BMM action may provide better diagnosis and treatment strategies for AML patients.

The BM microenvironment in AML is comprised of leukemia cells, stromal cells, endothelial cells and distinct immune cell subsets. Among them, vascular endothelial cells (ECs) promote leukemia cell proliferation, drug resistance, and recurrence through paracrine vascular endothelial growth factor (VEGF), adhesion, and fusion with leukemia cells, resulting in poor prognosis (Aguayo et al., 1999; Cogle et al., 2016; Padró et al., 2002; Wegiel, Ekberg, Talasila, Jalili, & Persson, 2009). Stromal cells can promote chemotherapy resistance in leukemia cells through ligand-receptor interactions (Hazlehurst & Dalton, 2001) such as SDF-1/CXCR4 (Tabe et al., 2007) and VLA-4/VCAM-1 (Jacamo et al., 2014). Adipocytes promote the proliferation, growth and chemotherapeutic resistance of leukemia cells by breaking down stored triglycerides into free fatty acids(Shafat et al., 2017) and secreting tumor-related proinflammatory cytokines (Ye et al., 2016).

The leukemia immune microenvironment presents with immune dysregulation and suppression, leading to an imbalance of suppressor T cells and effector T cells, T cell exhaustion and an increase in myeloid-derived suppressor cells (MDSCs) and leukemia-supporting macrophages compared to normal bone marrow tissue(Mendez, Posey, & Pandolfi, 2019). Recent studies on the characterization of the leukemia immune microenvironment could aid in the search for novel prognostic biomarkers and potential therapeutic targets (Vadakekolathu et al., 2020; Yan et al., 2019). In addition, treatments such as chemotherapy, immunotherapy, and combination therapy to alter the immune microenvironment of AML have been widely used (Allie, Zhang, Tsai, Noelle, & Usherwood, 2013; Assi, Kantarjian, Ravandi, & Daver, 2018). However, different immune cells have different effects in AML. Therefore, understanding the distribution and function of immune-related genes in different immune cells is of great significance to further explore the BM immune microenvironment of AML patients.

Here, we investigated the impact and potential mechanisms of immune-related genes on the prognosis of AML. We are the first group to screen for hub genes in this disease using multiomics analysis. We constructed a LASSO-Cox regression model to predict the prognosis of AML according to the characterization of the leukemia immune microenvironment. The distribution of hub genes in immune cells was revealed through single-cell sequencing data and may provide the potential for precise patient stratification and treatment.

## Materials and Methods

### Datasets

The test cohort of acute myelocytic leukemia (AML) was downloaded from The Cancer Genome Atlas (TCGA) database (https://www.cancer.gov/) and includes mRNA data from 151 cases (RNASeq V2), miRNA data from188 cases (Illumina HiSeq miRNAseq) and Illumina Human Methylation450 Bead Array data from 140 cases. Samples were selected for the study according to the following criteria: 1) acute myelocytic leukemia was pathologically diagnosed, 2) all three kinds of data were available for the patient, and 3) the clinical information was complete. Finally, 97 patients were selected for our following study.

The other datasets were obtained from the Gene Expression Omnibus (GEO) database (https://www.ncbi.nlm.nih.gov/geo/). A total of 4 AML patient datasets were employed as independent validation cohorts: GSE12417, containing 2 datasets in which 163 samples were analyzed using the GPL96 (Affymetrix Human Genome U133A Array) platform and 80 samples were analyzed using GPL570 (Affymetrix Human Genome U133 Plus 2.0 Array) platform; the GSE106291 dataset (251 samples), which was generated using the GPL18460 (Illumina HiSeq 1500, *Homo sapiens*) platform; and the GSE37642 dataset (137 samples), which was generated using the GPL570 (Affymetrix Human Genome U133 Plus 2.0 Array) platform.

The single-cell RNA sequence dataset GSE116256, including 16 untreated samples (D0), was used to reveal the distribution of hub genes in immune cell types. The immune gene set, including 776 genes, was acquired from a previous study (Charoentong et al., 2017).

### Screening of candidate genes and hierarchical clustering

Differential mRNA and miRNA expression were analyzed by the DESeq2 (Love, Huber, & Anders, 2014) function (*P*<0.05, |logFC|>1). For each probe in the methylation data, the value shown is the β value (β =U/(M+U+1)), where M is the methylated probe signal strength and U is the unmethylated probe signal value. The methylmix package (Gevaert, 2015) was used to analyze the correlation between the gene methylation level and mRNA expression value (Pearson correlation coefficient test, R>0.5, *P*<0.05). Unsupervised hierarchical clustering was performed (Euclidean distances and complete linkage method) based on SIGs to establish an immunogenomic classification of TCGA-AML patients.

### Immune infiltration analysis

The enrichment scores of 28 immune signatures in each AML sample were quantified using single-sample gene set enrichment analysis (ssGSEA) (Subramanian et al., 2005). Stromal, immune, and estimate scores were further calculated to evaluate tumor purity and immune cell infiltration in tumor tissues based on the mRNA expression data using the ESTIMATE (Estimation of STromal and Immune cells in MAlignant Tumor tissues using Expression data) algorithm (Yoshihara et al., 2013).

### Protein–protein interaction (PPI) network construction and gene ontology (GO) functional enrichment analysis

mRNA interaction data were obtained from the STRING database (https://string-db.org/) (Szklarczyk et al., 2019). The PPI network was established using Cytoscape software (Shannon et al., 2003). The Database for Annotation, Visualization and Integrated Discovery (DAVID, https://david.ncifcrf.gov/) (Huang da, Sherman, & Lempicki, 2009a, 2009b) was used for GO enrichment analysis (P < 0.05), and the results were plotted by the GO plot package (Walter, Sánchez-Cabo, & Ricote, 2015).

### Survival analysis and prognostic model construction

A Cox regression model was constructed to identify immune-related genes that significantly correlated with the OS (SIGs) of AML patients in TCGA. The genes that met the standard of *P* < 0.05 were used for subsequent study. Least absolute shrinkage selection operator (LASSO)-penalized Cox regression analysis (Friedman, Hastie, & Tibshirani, 2010; Simon, Friedman, Hastie, & Tibshirani, 2011) was applied to distinguish the most important SIGs and to construct a prognostic model for AML based on a linear combination of the regression coefficients. Kaplan-Meier survival curves and receiver operating characteristic (ROC) curves were used to test and validate the performance of the classifier.

### scRNA dataset analysis

We employed the Seurat (Butler, Hoffman, Smibert, Papalexi, & Satija, 2018; Stuart et al., 2019) and SingleR (Aran et al., 2019) packages to generate Uniform Manifold Approximation and Projection (UMAP) plots and reveal the distribution of hub genes in each immune cell type.

### Patients and samples

A total of 55 patients with newly diagnosed AML were enrolled between Jan 2016 to Dec 2020 at the Xinqiao Hospital of the Army Medical University in China. Samples were selected for the study according to the following criteria: 1) acute myelocytic leukemia diagnoses were based on morphological findings, karyotype, and immunophenotypic features of leukemia cells by consultant hematologist, 2) patients were newly diagnosed, and remained untreated at the time of collection, 3) the clinical information was complete, and 4) a total of 38 genes mutation and SNP were detected using next-generation sequencing (NGS) technology.

### Molecular Docking

The virtual screening of molecular docking was performed using AutoDock Vina 1.1.2 (Trott & Olson, 2010 Jan 30) to predict the most likely optimal ligands. The three-dimensional structure of CD52 (PDB ID: 6OBD), CLCN5 (PDB ID: 2J9L) and ICAM3 (PDB ID: 1T0P) were retrieved from the Protein Data Bank (https://www.rcsb.org/). A library of 2115 FDA approved compounds were extracted from ZINC15 druglike database (http://zinc.docking.org/). The visualization of active interactions between proteins and compounds was performed by Discovery Studio Visualizer v4.5.0 (BIOVIA).

## Results

### Classification of AML based on immune-related genes that significantly affect patient prognosis

For a more extensive study of immune genes in AML, we retrieved transcriptome, microRNA, and DNA methylation profile data and integrated clinical information for 97 samples from TCGA database (**Table S1**). A Cox proportional hazard regression model was employed to analyze 776 immune-related genes (Charoentong et al., 2017) in the mRNA expression data of 97 samples (*P*<0.05), and 98 survival-related immune genes (SIGs) that significantly affected the survival of AML patients were identified (**Table S2**).

Using unsupervised clustering analysis (Euclidean statistics and complete linkage method) of 98 SIGs, those 97 samples were clustered into three distinct immune subtypes (Im1: immune cluster 1, Im2: immune cluster 2, Im3: immune cluster 3) based on the gene expression signature (**Figure 1A**). As shown in immune gene heatmaps, most of the SIGs were highly expressed in the Im1 and Im3 clusters but expressed at low levels in the Im2 cluster (**Figure 1B**). Kaplan-Meier survival analysis revealed that the prognosis of the Im2 cohort was significantly better than that of the Im1 and Im3 cohorts (**Figure 1C,** log-rank test, *P*=0.0008).

**Figure 1.**
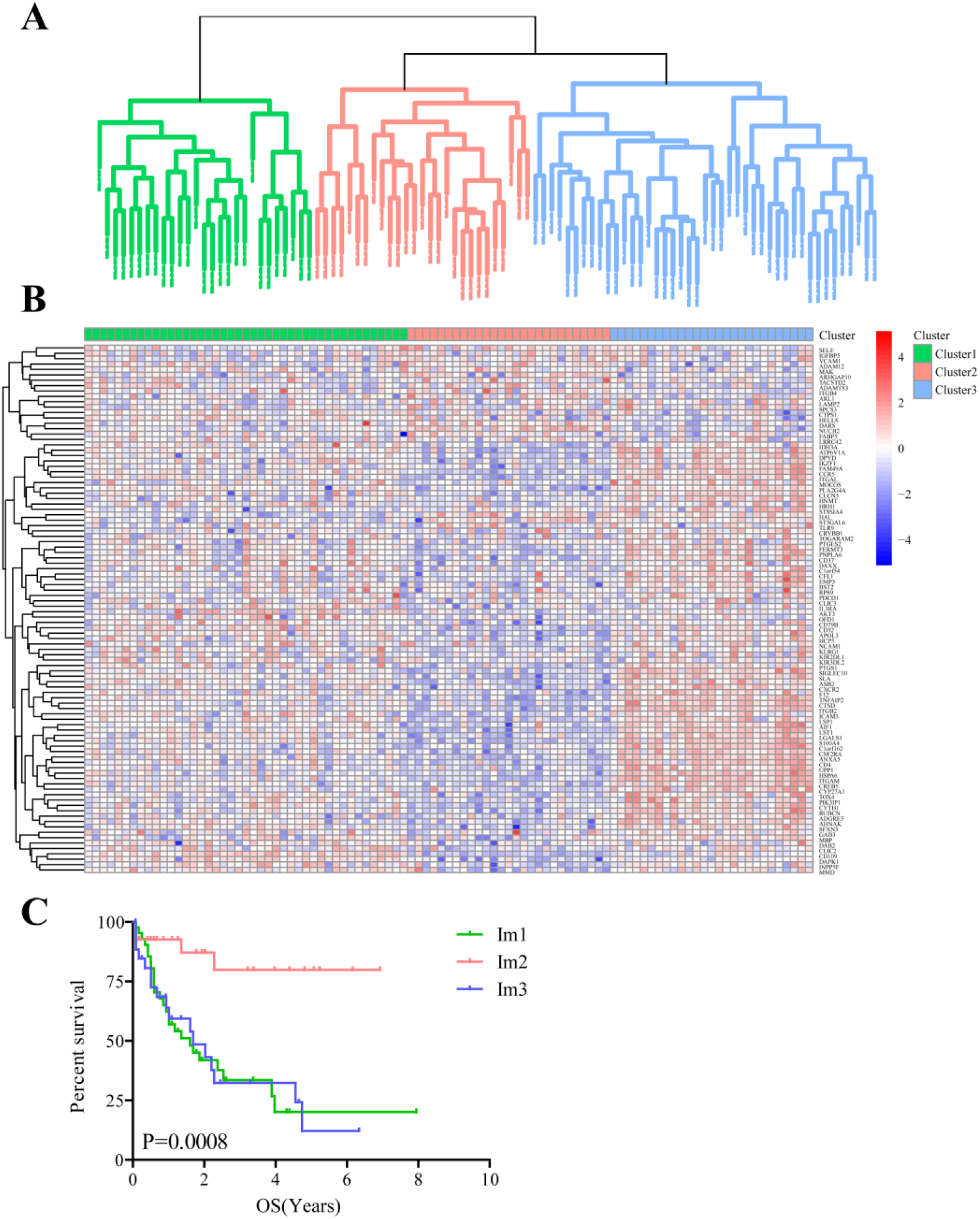
Unsupervised clustering analysis of AML patients based on 98 survival-related immune genes. **A.** All 97 TCGA-AML patients were divided into 3 clusters (green: Im1 cluster, red: Im2 cluster, blue: Im3 cluster). **B.** Heatmap of 98 survival-related immune genes in different AML clusters. **C.** The Kaplan-Meier survival analyses along with the Log-rank test were used to compare the overall survival (OS) of the Im1, Im2 and Im3 clusters.

### The immune infiltration was significantly different in different cluster

As the immune microenvironment was significantly correlated with the occurrence and development of AML, a single-sample gene set enrichment (ssGSEA) algorithm was utilized to explore differences in the immune microenvironment among the three immune clusters. The results showed that the Im2 cluster had fewer infiltrating immune cells than the Im1 and Im3 clusters (**Figure 2A**), and the specific infiltration of immune cells in different clusters is shown in **Figure S1**. Consistent findings demonstrated that the immune scores were significantly lower in the Im2 cluster (**Figure 2B**, unpaired t test, *P* < 0.001), while tumor purity was significantly higher in the Im2 cluster but significantly lower in the Im1 and Im3 clusters (**Figure 2C**, unpaired t test, *P* < 0.001). Generally, we can conclude that patients with less immune infiltration and lower immune scores may have a better prognosis than those with more immune infiltration and higher immune scores.

**Figure 2.**
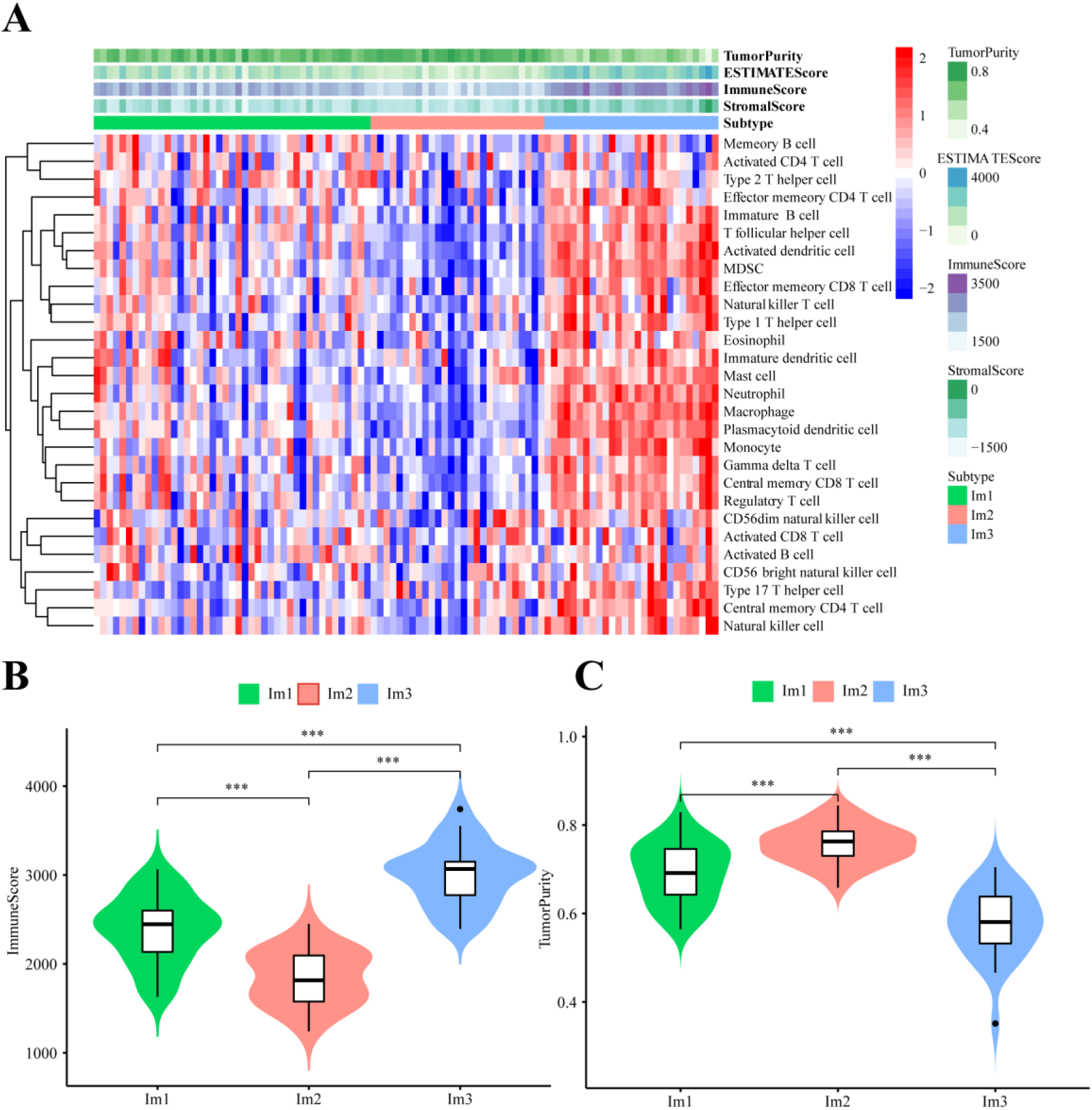
Immune functional characters of 3 AML patients clusters. **A.** Heatmap of the Im1, Im2 and Im3 cohorts of 97 TCGA-AML patients using ssGSEA scores from 28 immune cell types. Violin plots depict the immune score (**B**) and tumor purity (**C**) of the Im1, Im2 and Im3 cohorts.

### 42 DEG-SIGs were screened by mRNA expression data analyzing

Based on the significant differences in immune infiltration and survival trends between the Im2 cluster and Im1/3 cluster, we defined Im2 as the immune infiltration-lacking subtype (IL type) and Im1/3 as the immune infiltration-rich subtype (IR type). To reveal the potential mechanisms of different prognoses between IL and IR subtypes, an elaborate analysis of the mRNA expression profiles of the two types of AML patients was implemented. We performed differentially expressed gene analyses and identified 1936 differentially expressed genes (DEGs) with significant differences between IL and IR subtypes. There were 42 SIG-DEGs which were common members of 1936 DEGs and 98 SIGs (*P* < 0.05, |Fold Change|≥2) (**Table 1**) (**Figure 3A, B**).

**Table 1.** Common 42 intersecting genes of differentially expressed genes (DEGs) and survival-related immune genes (SIGs) between immune infiltration-lacking subtype (IL type) and immune infiltration-rich subtype (IR type) (P < 0.05, | Fold Change |≥2).

| DEG-SIGs | Parametric p-value | Fold-change |
| --- | --- | --- |
| ADAMTS3 | 0.022973 | 0.37 |
| AKT3 | 0.0002581 | 2.31 |
| ANXA5 | 0.00000001 | 5.6 |
| APOL3 | 0.00000009 | 3.11 |
| ASB2 | 0.00000017 | 3.41 |
| CCR5 | 0.00000025 | 5.73 |
| CD109 | 0.00000033 | 18.68 |
| CD4 | 0.0000012 | 3.11 |
| CD52 | 0.00000041 | 4.64 |
| CLCN5 | 0.0000017 | 2.78 |
| CLIC2 | 0.0003049 | 2.5 |
| CREB5 | 0.0000003 | 4.64 |
| CSF2RA | 0.00000049 | 2.82 |
| CTSD | 0.0000022 | 2.38 |
| CXCR2 | 0.0000106 | 3.13 |
| CYP27A1 | 0.0001964 | 4.2 |
| DAPK1 | 0.00000057 | 3.52 |
| DPYD | 0.00000065 | 2.97 |
| F12 | 0.0000034 | 2.91 |
| FAM49A | 0.0000331 | 2.5 |
| HAL | 0.0014438 | 0.44 |
| HCP5 | 0.0000109 | 2.31 |
| HNMT | 0.045659 | 2.23 |
| HRH1 | 0.0082122 | 2.03 |
| HSPA6 | 0.0000058 | 3.28 |
| ICAM3 | 0.0000006 | 2.08 |
| ITGAL | 0.00000073 | 3.37 |
| ITGAM | 0.00000081 | 4.61 |
| ITGB2 | 0.00000089 | 3.93 |
| LGALS1 | 0.0000022 | 3.18 |
| LSP1 | 0.00000097 | 4.77 |
| LST1 | 0.00000105 | 5.03 |
| NUCB2 | 0.00000113 | 0.43 |
| PLA2G4A | 0.00000121 | 3.87 |
| PTGS1 | 0.0000015 | 2.24 |
| S100A4 | 0.00000129 | 3.65 |
| SELE | 0.0140379 | 0.48 |
| SFXN3 | 0.00000137 | 2.66 |
| SIGLEC10 | 0.0000046 | 2.53 |
| SLA | 0.00000145 | 3.17 |
| TNFAIP2 | 0.00000153 | 5.44 |
| UPP1 | 0.000643 | 2.29 |

**Figure 3.**
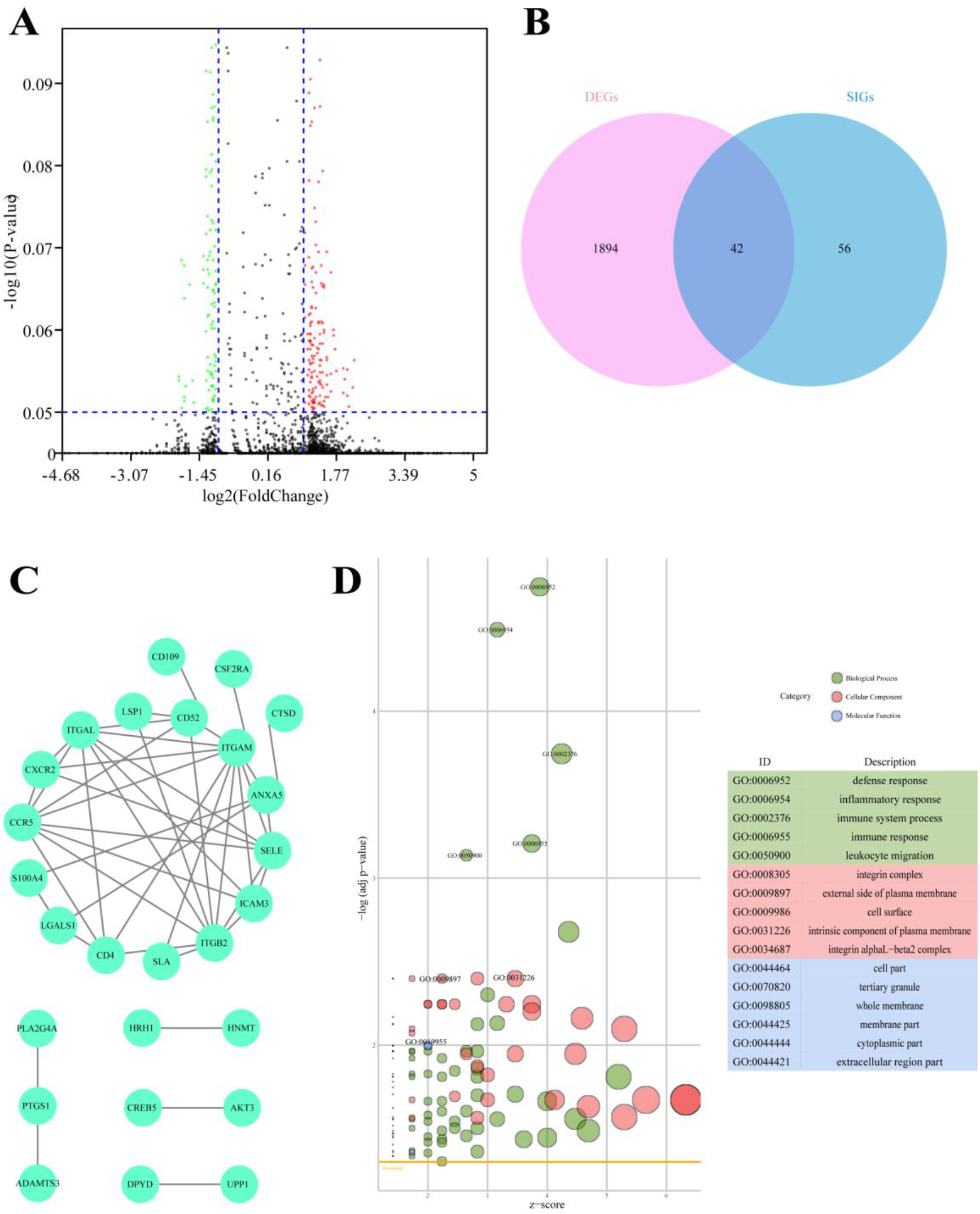
Difference analysis of the mRNA expression dataset from TCGA-AML patients. **A.** Volcano plot of differentially expressed genes between immune infiltration-lacking subtype (IL type) and immune infiltration-rich subtype (IR type). **B.** Venn plot of the intersecting genes between differentially expressed genes (DEGs) and survival-related immune genes (SIGs). **C.** PPI network of 42 overlap genes DEG-SIGs. **D.** The bubble plot represents the GO functional enrichment analysis of 42 DEG-SIGs.

To elucidate the mechanism of prognosis difference between IL and IR subtype we analyzed the interaction on the STRING website (https://string-db.org/) and constructed protein-protein interaction (PPI) networks to draw visualized interactome maps of the 42 DEG-SIGs (**Figure 3C**). Gene ontology (GO) functional enrichment analysis distinguished some enriched terms in three subontologies: biological processes (BP), cellular component (CC), and molecular function (MF) (**Figure 3D**). For BP, 42 DEG-SIGs were enriched in defense response, inflammatory response, and immune system process. With regard to CC, 42 DEG-SIGs were enriched in integrin complex, external side of plasma membrane, and cell surface. For MF, 42 DEG-SIGs were enriched in cell part, tertiary granule, and whole membrane. These results may partially illustrate the potential mechanisms of 42 DEG-SIGs affecting the prognosis of AML patients.

### 19 hub genes were screened by integrated analysis of mRNA expression data, miRNA expression data and methylation data

Considering the complex mechanism of leukemogenesis and progression, we next conducted integrated multiomics analysis to identify hub genes that were associated with prognosis. Comparing the miRNA expression profiles of patients between IL and IR subtypes, we revealed 93 miRNAs that were significantly differentially expressed (*P*<0.05, |FC|≥2) (**Figure 4A**). A total of 7294 target miRNA genes (TDEmiRs) were identified using DIANO TOOLS/microT-CDS (threshold=0.9). Through integrated bioinformatics analysis, we selected 15 commonly differentially expressed genes from 42 DEG-SIGs and 7294 TDEmiRs between the IL and IR subtypes (**Figure 4C**).

**Figure 4.**
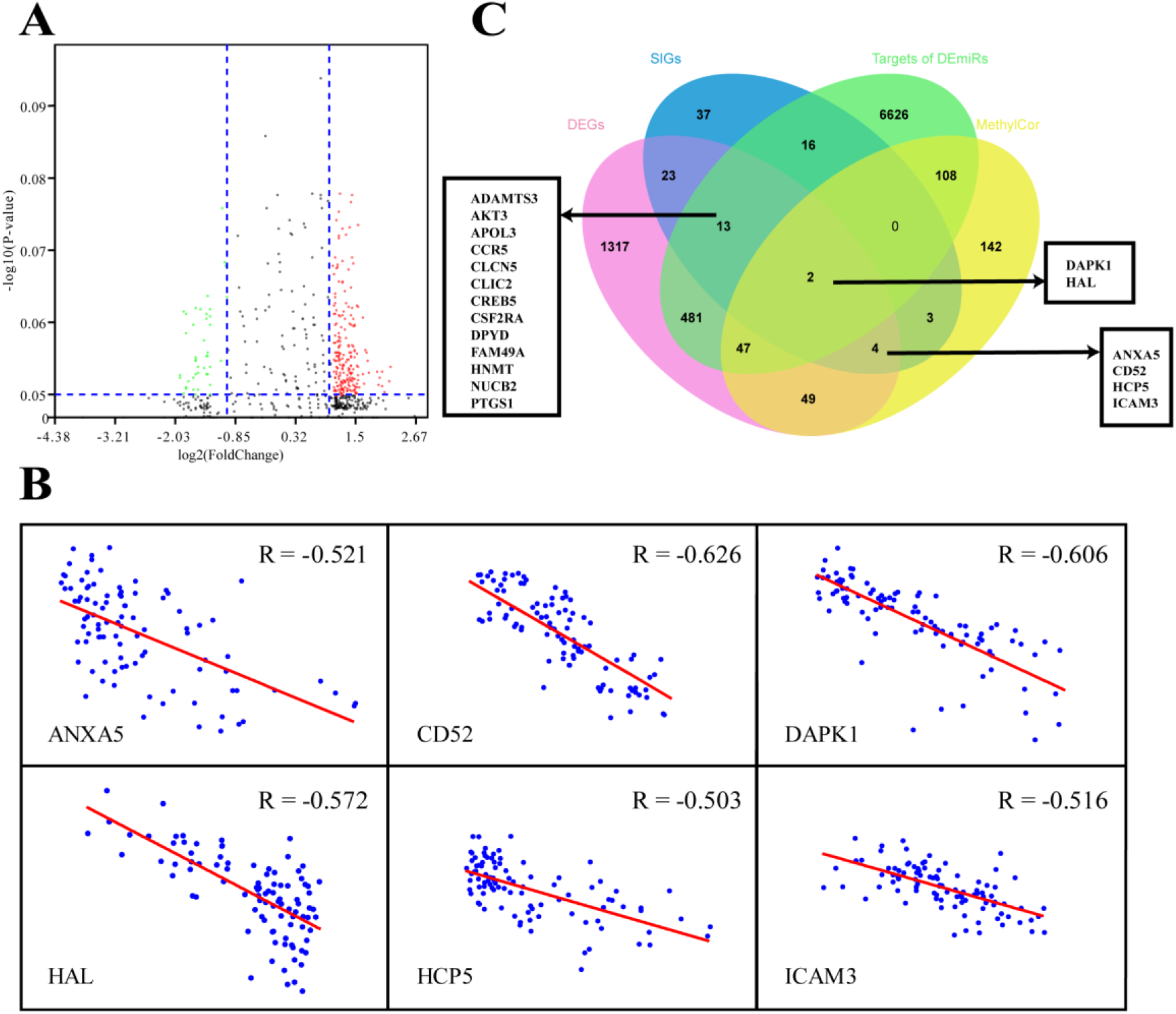
Multiomics analysis of 97 TCGA-AML patients. **A.** Volcano plot of differentially expressed miRNAs between IL and IR types. **B.** Correlation between mRNA expression and DNA methylation level of 6 DEG & MethylCor genes. **C.** Venn plot of the intersection of DEGs, SIGs, targets of DEmiRs and MethylCor gene set.

Combined analysis of mRNA and methylation profiles indicated that there were significantly negative correlations between mRNA expression level and degree of methylation for 355 genes (R < - 0.5, p< 0.05). When these 355 methylation correlation genes (MethylCor) were cross-referenced with the 42 DEG-SIGs, we identified 6 common genes associated with immune infiltration and differential expression, methylation and prognosis between IL and IR subtypes (**Figure 4B, C**).

### A prognostic model based on 5 hub genes was constructed

Having observed significant differences in immune infiltration, gene expression and clinical behavior between IL and IR types, we next developed a LASSO-Cox proportional hazards regression model based on 19 immune-correlated DEGs by combining microRNA and epigenetic regulation data. Using the LASSO model, we built a classifier based on the 5 hub genes to predict the prognosis of AML (risk score = −0.086×ADAMTS3 + 0.180×CD52 + 0.472×CLCN5 - 0.356×HAL + 0.368×ICAM3) (**Figure 5A, B**). Kaplan-Meier plots displayed OS differences between patients in various subtypes (*P*=3.931×10^-06^) (**Figure 5C**), and the ROC curve suggested that the model can effectively predict the 1-, 3- and 5-year prognosis of AML (AUC=0.82, 0.83, 0.99, respectively) (**Figure 5D**). Consistent with earlier analysis, we found similar predictive performance for 151 mRNA samples of the TCGA-AML profile (*P*=3.369×10^-06^, AUC=0.63, 0.74, 0.83) (**Figure 5E, F**).

**Figure 5.**
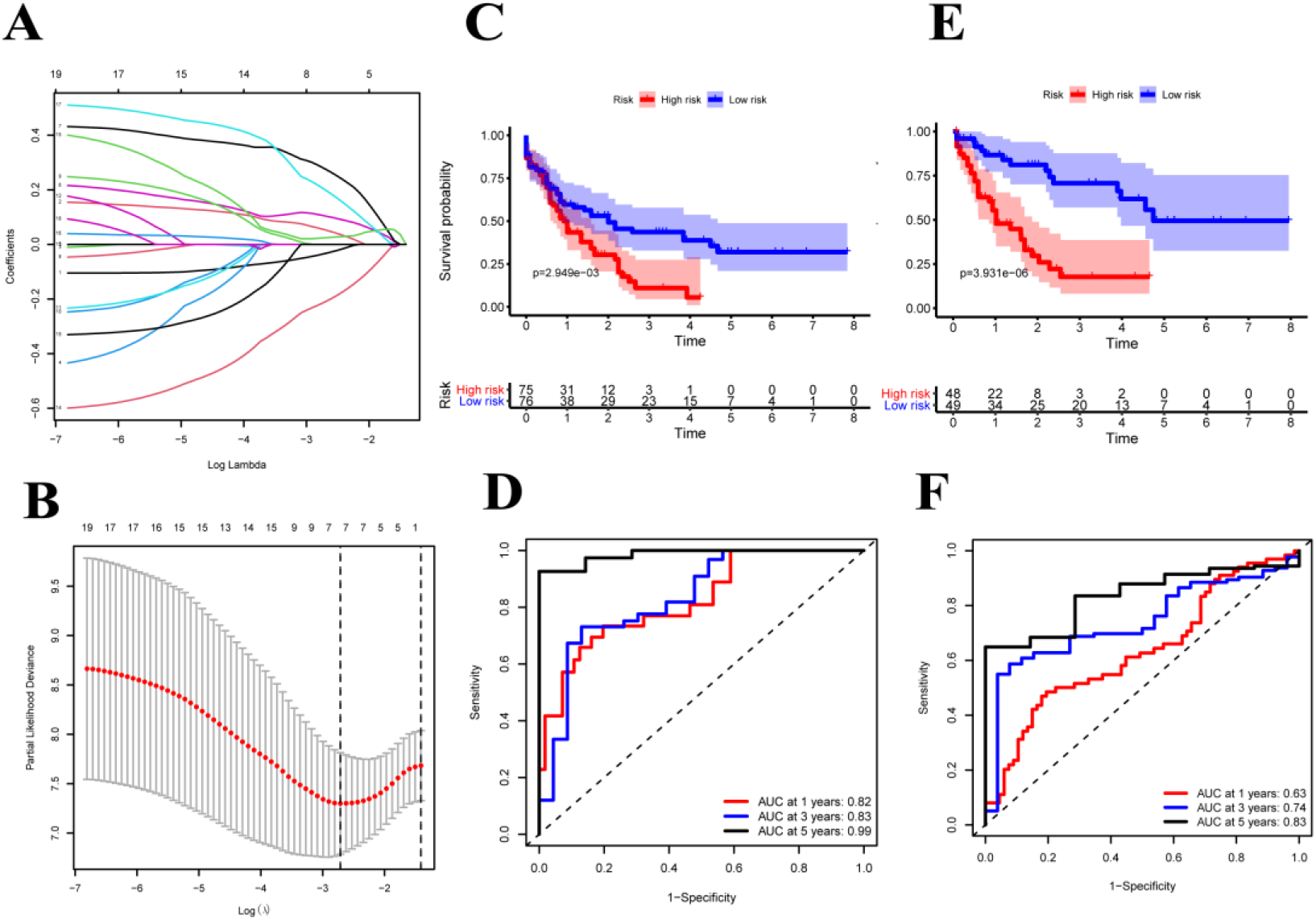
Construction of the COX regression model. **A.** LASSO coefficient profiles of 19 candidate SIGs. **B.** Tuning parameter (λ) selection cross-validation error curve. The vertical lines were drawn at the optimal values determined by the minimum criteria and the 1-SE criteria. **C, E.** OS in patients with high vs. low risk scores depicted by Kaplan-Meier plots in the TCGA-AML-97 and TCGA-AML-151 cohorts. **D, F.** ROC curves depicting the accuracy of the Cox regression model in identifying AML subtypes with poor prognosis in the TCGA-AML-97 and TCGA-AML-151 cohorts.

### The prognostic model can still predict the prognosis of AML patients effectively in various validation cohorts

Since APL can achieve good therapeutic effects with the application of ATRA and ATO, we performed predictive tests separately in APL and non-APL patients. Surprisingly, we observed the same conclusions in APL (log-rank test, *P*=0.3618, AUC=0.85, 0.88, 0.78) and non-APL (log-rank test, *P*=0.01396, AUC=0.6, 0.68, 0.82) (**Figure 6A, B**). To test this model further, validation cohorts were obtained from the GEO database. Kaplan-Meier plots and ROC curves at 1, 3 and 5 years confirmed the prognostic value of the 5-hub-gene-based model: GSE106291 (log-rank test, *P*=3.311×10^-06^, AUC=0.66, 0.66, 0.61), GSE12417-GPL96 (log-rank test, *P*=3.377×10^-02^, AUC=0.61, 0.6, NA), GSE12417-GPL570 (log-rank test, *P*=3.763×10^-02^, AUC=0.56, 0.64, NA), and GSE37642 (log-rank test, *P*=3.966×10^-02^, AUC=0.51, 0.57, 0.6) (**Figure 6C, D, E, F**). After stratification by disease classification, the results showed that the risk score of the IL type was significantly lower than that of the IR type (**Figure S2**). These evaluations demonstrated that the 5-hub-gene-based model could identify a group of high-risk patients within conventionally assigned risk groups and may guide clinical practice.

**Figure 6.**
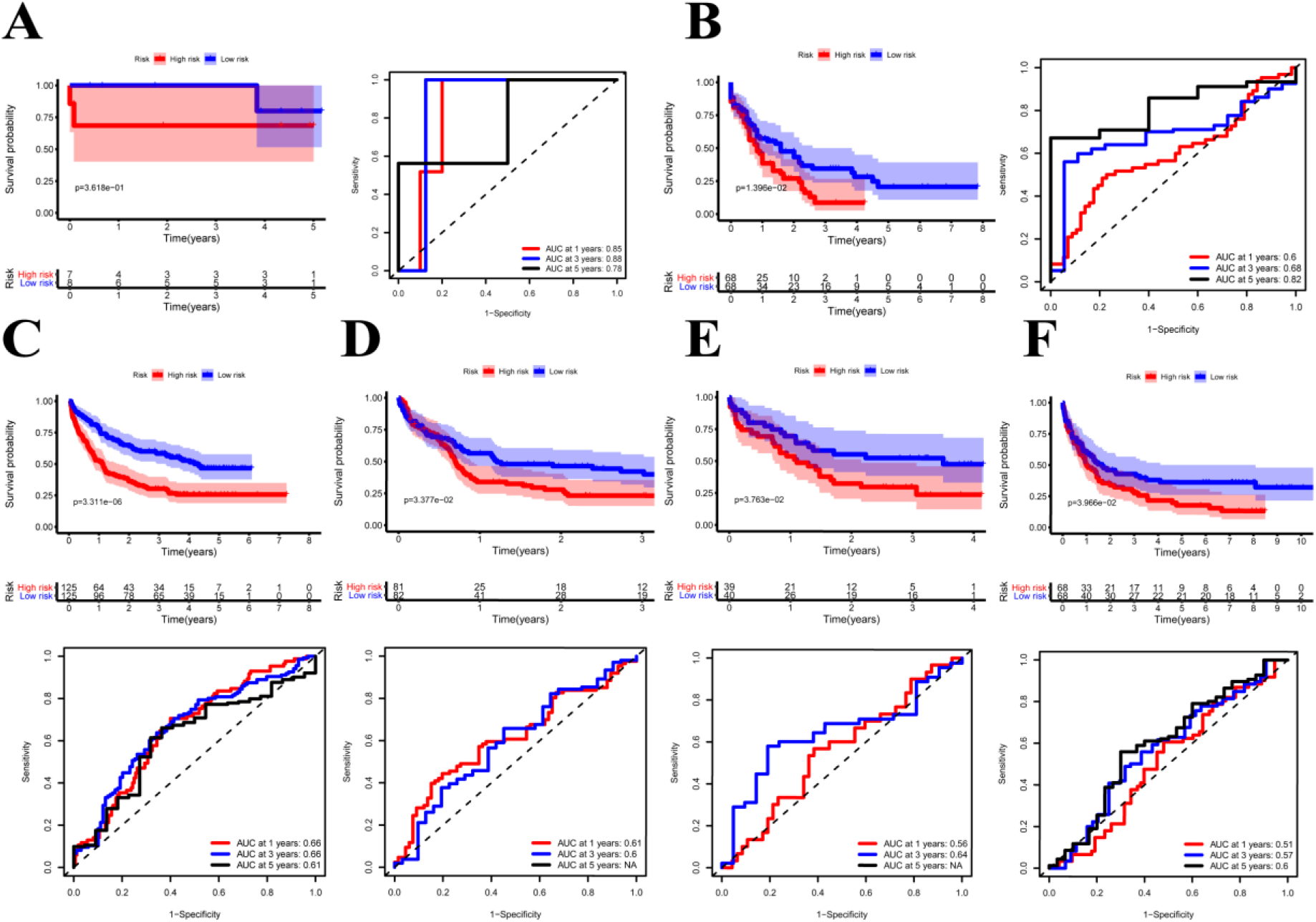
Validation of the prognostic model. **A, B.** Kaplan-Meier plots and ROC curves of TCGA-AML-APL and TCGA-AML-nonAPL cohorts. **C, D, E, F.** Kaplan-Meier plots and ROC curves of 3 GEO cohorts.

### The better efficacy of the prognostic model for the prognosis of AML patients was further verified by clinical samples

For verifying the prognostic value in the 5-hub-gene-based model, we collected 6575 genes mutation detected in 200 newly diagnosed AML patients (TCGA.LAML.mutect.somatic.maf, https://portal.gdc.cancer.gov/files/27f42413-6d8f-401f-9d07-d019def8939e) and 38 genes mutation detected in 55 newly diagnosed AML patients (Xinqiao Hostpital). The common mutated genes were *DNMT3A*, *IDH1*, *NRAS*, *RUNX1* and *TET2*. In this model classification, the high risk was significantly associated with the mutation of *RUNX1* (p=0.015) and *TET2* (p=0.054) considered by chi-square test (**Table 2**). Kaplan-Meier analyses of 55 patients with prognostic information indicated that patients with mutation of *RUNX1* (**Figure 7A**, p=0.0001) and *TET2* (**Figure 7B**, p=0.2257) were correlated with a poor prognosis and had a shorter median survival duration.

**Table 2.** Results of Chi-square test to 5 common mutated genes based on regrouping LASSO model in TCGA-AML database.

|  | Regroup | Mutated sample size | Normal sample size | P-value |
| --- | --- | --- | --- | --- |
| *DNMT3A* | high | 12 | 40 | 0.326 |
|  | low | 8 | 46 |  |
| *IDH1* | high | 4 | 48 | 0.201 |
|  | low | 1 | 53 |  |
| *NRAS* | high | 3 | 49 | 0.358 |
|  | low | 1 | 53 |  |
| *RUNX1* | high | 8 | 44 | 0.015 |
|  | low | 1 | 53 |  |
| *TET2* | high | 4 | 48 | 0.054 |
|  | low | 0 | 54 |  |

**Figure 7.**
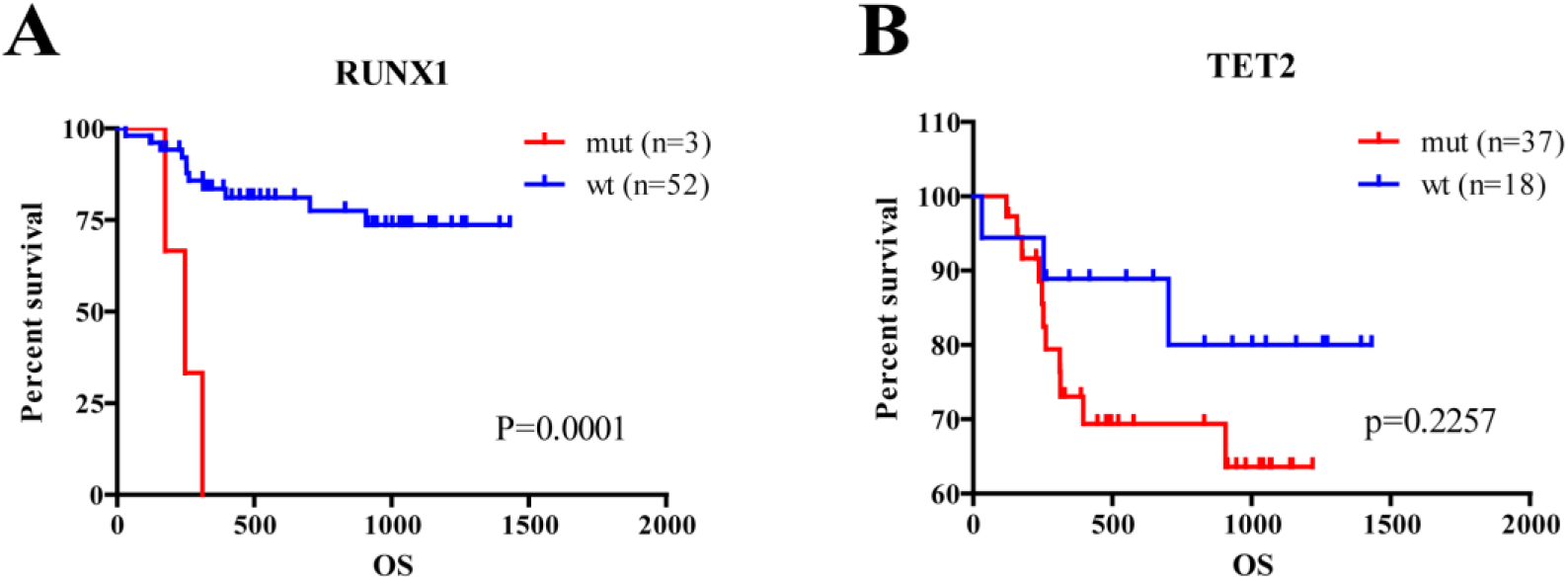
Analysis of hub genes and mutated genes in AML. **A.** Kaplan-Meier estimates of the OS according to *RUNX1* mutation status. **B.** Kaplan-Meier estimates of the OS according to *TET2* mutation status. (wt: wild type, mut: mutation)

### The diverse distribution of hub genes in immune cells of AML patients

To explore the value of these five hub genes in AML pathogenesis, we further identified the single-cell sequencing dataset GSE116256 to describe the distribution of the 5 hub genes in immune cells using the Seurat package for clustering and the SingleR package for annotation (**Figure 8A**). As shown in the scatter plot (**Figure 8B**) and violin plot (**Figure 8C**), CD52, ICAM3 and CLCN5 were widely expressed in granulocytes, monocytes, T lymphocytes, B lymphocytes, dendritic cells and NK cells, whereas ADAMTS3 was rarely expressed in those cells. HAL is highly expressed in granulocytes and monocytes but rarely expressed in other immune cells. Accordingly, we hypothesized that these hub genes play various roles through the regulation of gene expression in specific cells. The hub gene expression of blood cells in the Protein Atlas database (https://www.proteinatlas.org/) further confirmed this result (**Figure S3**).

**Figure 8.**
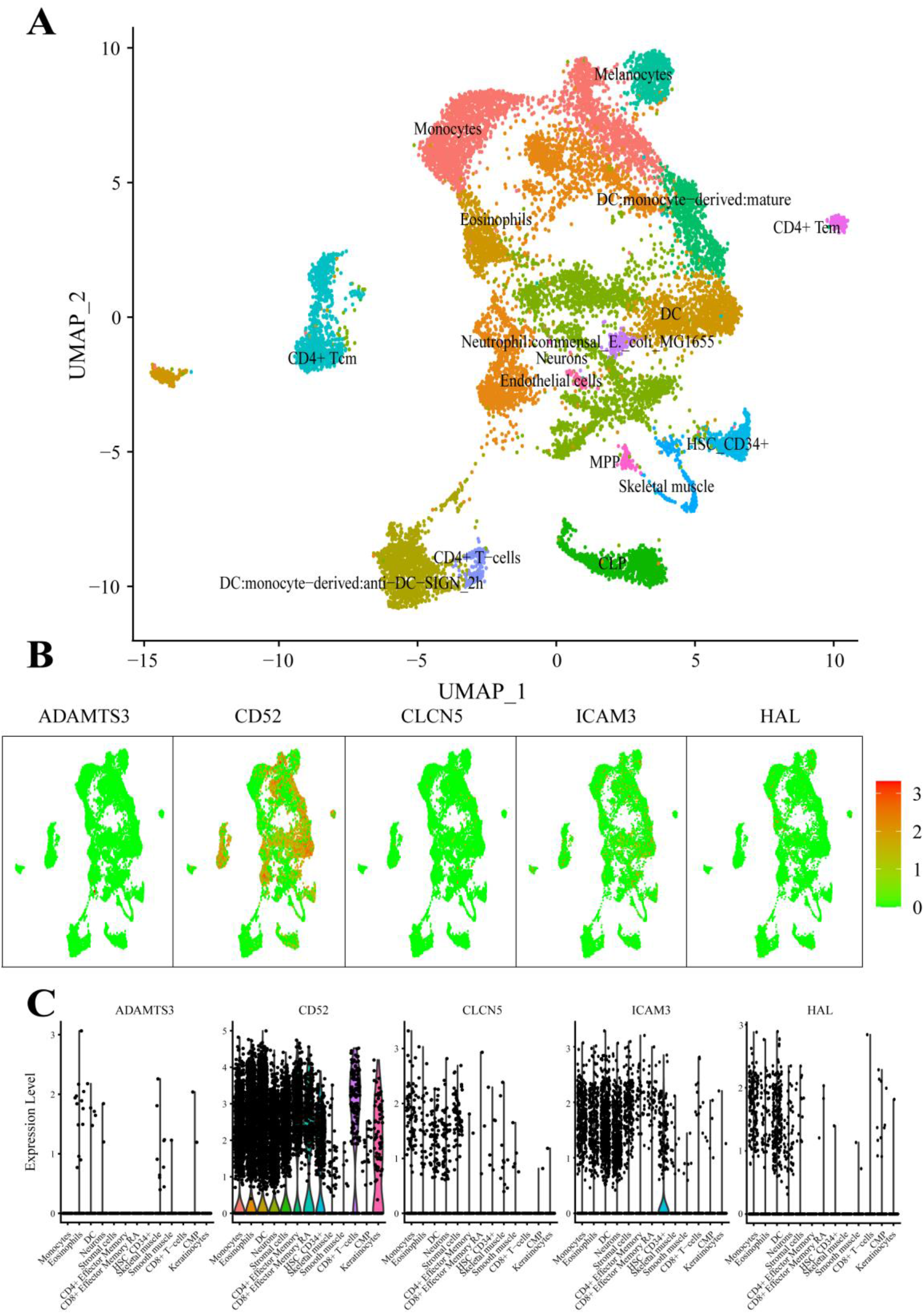
scRNA analysis of hub genes. **A.** Clustering analysis of the UMAP plot, color coded based on cell types. **B.** Overlaying gene expression on UMAP clusters to illustrate the distribution of hub genes in each cell type. **C.** Violin plots of hub genes in each cell type.

### Investigation of best-fitting compounds on hub genes

To investigate best-fitting compounds, we performed virtual screening of molecular docking using the three-dimensional structure of CD52 (PDB ID: 6OBD), CLCN5(PDB ID: 2J9L), ICAM3(PDB ID: 1T0P) and 2115 FDA approved compounds in ZINC15 database. The predicted binding affinities of the top 2 hit compounds against respective targets are ranked from highest to lowest. Binding energy (Kcal/mol) for interaction of proteins and compounds are as follows: CD52 with ZINC164528615 (Glecaprevir) (−6.4 Kcal/mol), CD52 with ZINC3938684 (Toposar) (−6.3 Kcal/mol); ICAM3 with ZINC52955754 (Ergotamine) (−8.3 Kcal/mol), ICAM3 with ZINC1612996 (Irinotecan) (−8.2 Kcal/mol); CLCN5 with ZINC3978005 (Dihydroergotamine) (−11.8 Kcal/mol), CLCN5 with ZINC52955754 (Ergotamine) (−11.5 Kcal/mol). 2D visualization of the most probable interactions of these proteins and candidate compounds are represented in **Figure 9.**

**Figure 9.**
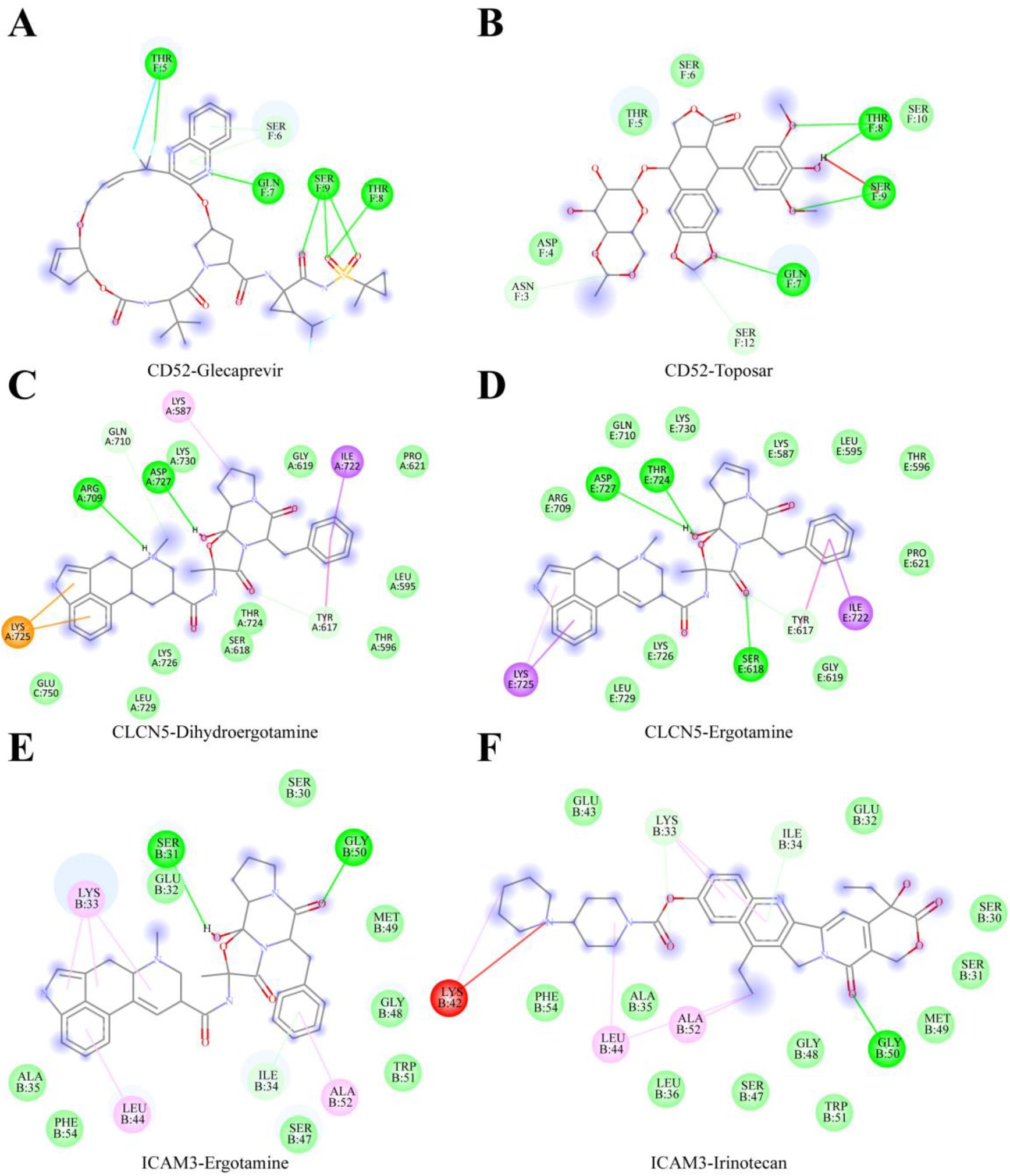
Shows 2D interaction representations of the best pose of **A.** CD52 with ZINC164528615 (Glecaprevir), **B.** CD52 with ZINC3938684 (Toposar); **C.** ICAM3 with ZINC52955754 (Ergotamine), **D.** ICAM3 with ZINC1612996 (Irinotecan); **E.** CLCN5 with ZINC3978005 (Dihydroergotamine), **F.** CLCN5 with ZINC52955754 (Ergotamine).

## Discussion

AML is a highly heterogeneous malignant tumor with poor prognosis. A large number of studies have demonstrated that the immune microenvironment of AML patients is altered significantly to promote leukemogenesis in AML(Mittal, Gubin, Schreiber, & Smyth, 2014; Teague & Kline, 2013). In this study, we classified AML patients according to the difference in the SIG expression signature. The results showed that the prognosis of AML patients with immune infiltration deficiency (IL subtype) was significantly better than that of patients with immune infiltration enrichment (IR subtype), which was contrary to the effect of the immune infiltration degree in solid tumors (Thorsson et al., 2018). Recent studies have shown that the heterogeneity of immune infiltration may be affected by the components of cytokines in the microenvironment(Benci et al., 2016; J. Li et al., 2018) and the expression of tumor driver genes(Bezzi et al., 2018). These heterogeneities were significantly associated with immunotherapeutic effects.

We further compared the gene expression profiles of IL and IR patients and found that the functions of genes with differential expression between the two subtypes were mainly enriched in defense response, inflammatory response and immune system process (BP); integrin complex, plasma membrane and cell surface (CC); and cellular partial, tertiary granules, and whole membrane (MF). Consistent with our results, several studies have confirmed that the biological functional diversity of the immune system is significantly altered in the BM immune microenvironment.

Previous studies have constructed many prognostic models on the basis of diverse omics data. *Hu* et al. constructed a DNA methylation-based prognostic model for AML patients(Hu et al., 2019), but the accuracy of the model (AUC=0.67-0.75) was lower than that of our model (AUC=0.82-0.99). A three-miRNA signature was built for non-M3 AML patients by *Xue* et al., but due to the lack of validation by any independent data, its clinical application value may need further verification (Xue et al., 2019). A four-gene-based prognostic model from Huang et al. (Huang, Liao, & Li, 2017) was also less accurate (AUC=0.66-0.71) than our model.

These conventional prognostic analyses generally consider only mRNA, miRNA, or methylation data. To the best of our knowledge, this is the only study to date to construct a prognostic model through the integrated analysis of multiple omics datasets (mRNA, miRNA and methylation data). The TCGA-AML test dataset and three independent GEO-AML validation datasets confirm that the model is very effective in predicting the prognosis of AML patients. The model consists of five hub genes: ADAMTS3, CD52, CLCN5, ICAM3 and HAL.

ADAMTS3 is a member of the ADAMTS family and plays an important role in the genesis and development of a variety of tumors(Gomis-Rüth, 2009; Rocks et al., 2008). Previous studies demonstrated that ADAMTS3 is expressed in mast cells(Charoentong et al., 2017), and mast cells have both tumor-promoting and tumor-inhibiting effects(Oldford & Marshall, 2015), thus leading to different prognoses in different tumors(Fleischmann et al., 2009; Groot Kormelink, Abudukelimu, & Redegeld, 2009; Rajput et al., 2008). Our study confirmed that ADAMTS3 expression was significantly negatively correlated with the prognosis of AML and is rarely expressed in other immune cells. Therefore, we will pursue further in-depth analysis of how ADAMTS3 exerts its antitumor effect through mast cells to explore its potential therapeutic value.

CLCN5 (chloride voltage gated channel 5) is a member of the chloride channel family. It is widely expressed in a variety of tumor cells(Ernest, Weaver, Van Duyn, & Sontheimer, 2005) and can enhance the chemotherapy resistance of chronic lymphocytic leukemia and multiple myeloma(Ruiz-Lafuente et al., 2015; Zhang et al., 2018). Hsa-let-7c-5p and hsa-mir-495-3p, two miRNAs that target CLCN5, were found to be involved in the occurrence and development of a variety of tumors(Liao et al., 2020; Wu et al., 2020). Our study confirmed that CLCN5 plays a role in promoting tumorigenesis in AML, but its specific mechanism needs deeper investigation.

HAL (histidine ammonia lyase) is the rate-limiting enzyme of histidine catabolism(Brand Lm Fau - Harper & Harper, 1976). Kanarek et al. found that tumor cells with higher HAL expression levels possess higher sensitivity to methotrexate. Acute lymphocyte leukemia (ALL) patients with higher HAL expression were more likely to benefit from methotrexate treatment(Kanarek et al., 2018). In our study, we found that the expression level of HAL was significantly correlated with the degree of gene methylation. However, as a demethylation agent, hypomethylating agent (HMA) alone has difficulty achieving the intended effect in the clinical treatment of AML(Duchmann & Itzykson, 2019). We speculated that the combination of methotrexate and HMA may achieve better efficacy when changes in HAL expression are detected. Surprisingly, we found that HAL was only expressed in granulocytes and monocytes. Consistent with AML cells, granulocytes and monocytes originate from myeloid precursor cells(Mendez et al., 2019). Therefore, insight into the methylation mechanism of HAL and its effect on the innate immune response and AML cells will be conducive to improving the therapeutic effect of demethylation agents. We also assessed the interaction between HAL and hsa-miR-582-3p, which has been previously indicated as a tumor suppressor miRNA in AML (H. Li et al., 2019). Thus, deeper investigation of the relationship between HAL and miR-582-3p will be helpful to further understand the potential mechanism of malignant progression of AML.

Our study found that CD52 was widely expressed in leukemia cells and all types of immune cells. Evidence has revealed the high expression of CD52 in CD34+ stem cells of AML (5q-) patients and a significantly negative correlation between the CD52 expression value and the prognosis of AML (Blatt et al., 2014), which is consistent with our results. ICAM-3 (intercellular adhesion molecule 3, CD50) belongs to the ICAM (intercellular adhesion molecule) immunoglobulin superfamily and plays important roles in immune response and tumor development(Cavallaro & Christofori, 2004; Xiao, Mruk, & Cheng, 2013). Consistent with our scRNA sequencing results, ICAM-3 is expressed in granulocytes, monocytes and lymphocytes(de Fougerolles & Springer, 1992). In vivo and in vitro experiments confirmed that ICAM-3 participates in the proliferation, stemness and radiotherapy resistance of various tumors through the FAK pathway, PI3K/Akt pathway or other mechanisms(Chung et al., 2005; Kim et al., 2006; Shen et al., 2018). Interestingly, we found that the expression levels of CD52 and ICAM-3 were both significantly correlated with the degree of methylation but had opposite impacts on the prognosis of AML patients. Therefore, the roles of CD52 and ICAM-3 should be considered in HMA reagent treatment.

In conclusion, using a multiomics analysis and validation approach, we constructed and validated a novel, 5-hub-gene-based model that allows robust risk stratification and facilitates the identification of prognosis in AML. The distribution of the 5 hub genes in immune cells was revealed through scRNA sequencing analysis. Furthermore, we preliminarily explored the possible mechanisms of these hub genes in the leukemogenesis of AML. These findings may provide novel therapeutic targets for AML. Our study also provided new insight into the use of demethylation drugs in AML. Of course, this study is limited to bioinformatics analysis, and the proposed approaches need to be further tested in the clinic.

## Supplementary Data

**Figure S1.**
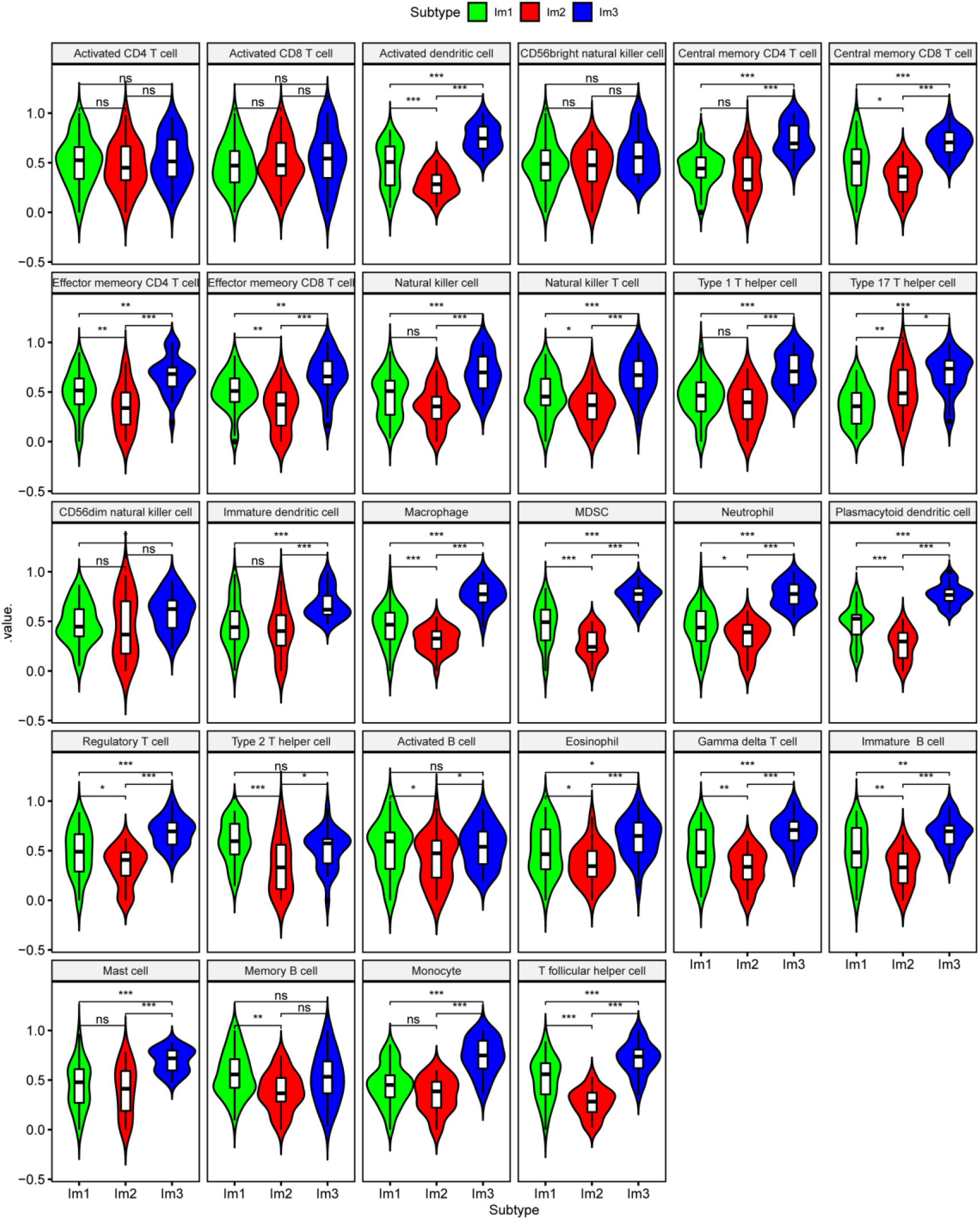
Comparison of the infiltration of immune cells in Im1, Im2 and Im3 clusters of AML patients.

**Figure S2.**
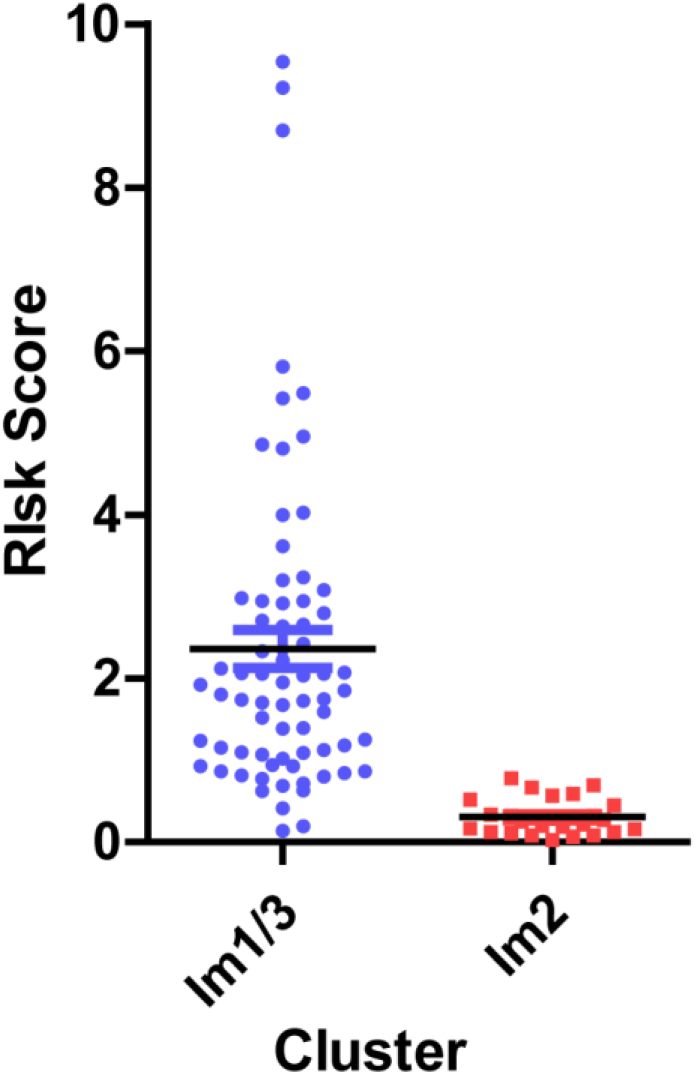
Comparison of the risk score between Im2 (immune infiltration-lacking subtype, IL type) and Im1/3(immune infiltration-rich subtype, IR type) clusters.

**Figure S3.**
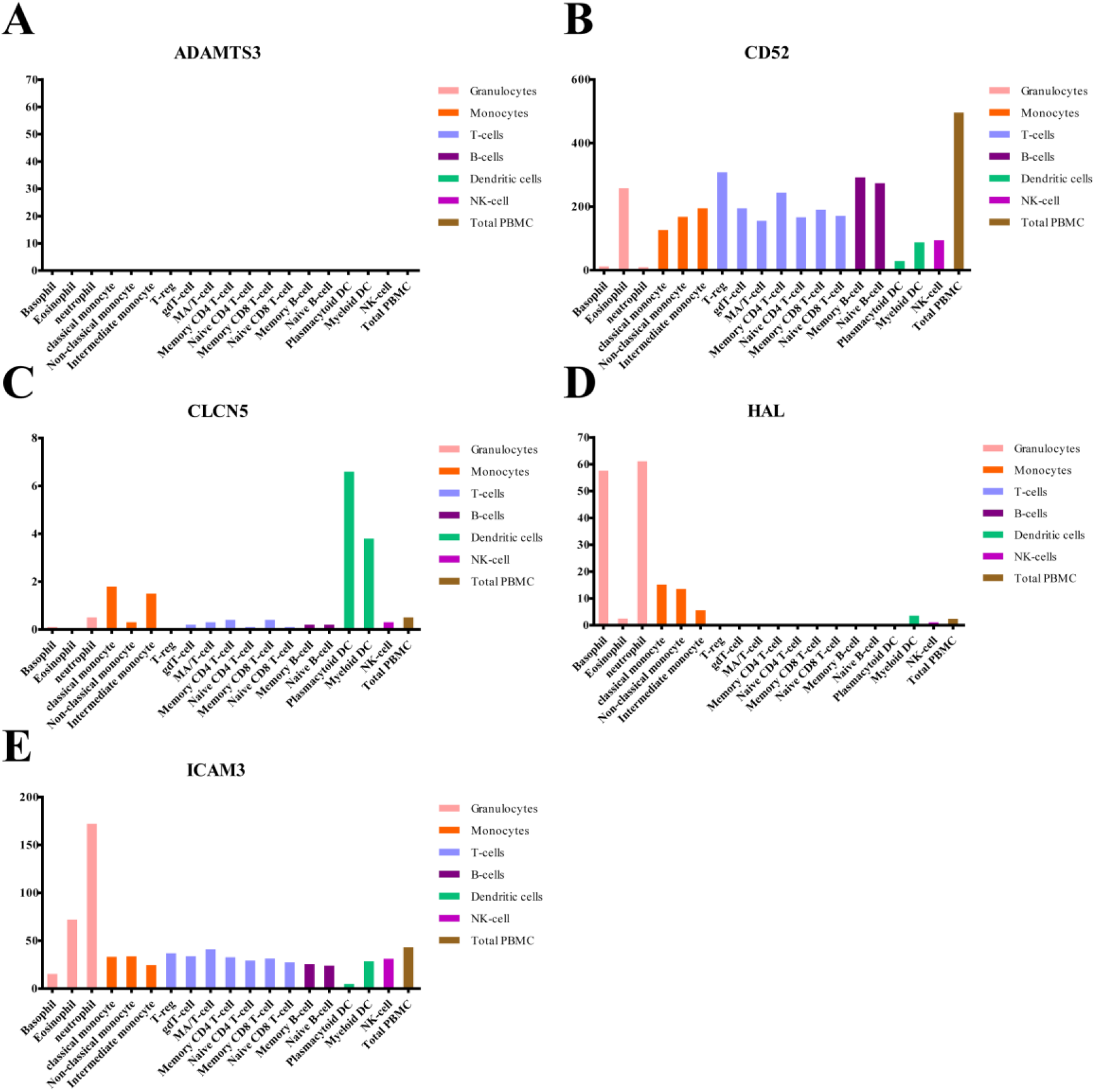
Histogram of hub gene expression in immune cells based on the Atlas database.

**Table S1.** Clinical characters of TCGA-AML patients.

| ID | Gender | Age | OS | Status |
| --- | --- | --- | --- | --- |
| TCGA-AB-2808 | male | 23 | 95.4 | alive |
| TCGA-AB-2819 | female | 52 | 83.2 | alive |
| TCGA-AB-2821 | male | 64 | 27.4 | dead |
| TCGA-AB-2828 | male | 55 | 76.1 | alive |
| TCGA-AB-2835 | male | 48 | 55.8 | alive |
| TCGA-AB-2846 | female | 57 | 46.7 | dead |
| TCGA-AB-2847 | male | 53 | 20.3 | dead |
| TCGA-AB-2849 | male | 39 | 74.0 | alive |
| TCGA-AB-2862 | female | 33 | 47.7 | alive |
| TCGA-AB-2863 | male | 63 | 1.0 | dead |
| TCGA-AB-2869 | female | 64 | 8.1 | alive |
| TCGA-AB-2870 | male | 76 | 5.1 | dead |
| TCGA-AB-2871 | male | 51 | 5.1 | alive |
| TCGA-AB-2872 | male | 42 | 21.3 | alive |
| TCGA-AB-2873 | female | 51 | 9.1 | alive |
| TCGA-AB-2874 | male | 59 | 13.2 | alive |
| TCGA-AB-2875 | male | 43 | 7.1 | alive |
| TCGA-AB-2876 | female | 45 | 40.6 | alive |
| TCGA-AB-2877 | female | 60 | 21.3 | alive |
| TCGA-AB-2878 | female | 47 | 12.2 | dead |
| TCGA-AB-2880 | male | 24 | 14.1 | dead |
| TCGA-AB-2881 | female | 48 | 13.1 | alive |
| TCGA-AB-2882 | female | 73 | 12.2 | dead |
| TCGA-AB-2883 | male | 60 | 24.3 | alive |
| TCGA-AB-2884 | female | 44 | 24.4 | dead |
| TCGA-AB-2885 | male | 71 | 7.1 | dead |
| TCGA-AB-2886 | male | 26 | 6.0 | alive |
| TCGA-AB-2888 | male | 57 | 11.1 | alive |
| TCGA-AB-2889 | male | 55 | 10.1 | alive |
| TCGA-AB-2892 | female | 42 | 31.4 | alive |
| TCGA-AB-2894 | female | 50 | 6.0 | dead |
| TCGA-AB-2895 | female | 41 | 5.1 | dead |
| TCGA-AB-2896 | female | 21 | 7.1 | dead |
| TCGA-AB-2897 | female | 50 | 8.1 | alive |
| TCGA-AB-2898 | female | 69 | 13.1 | alive |
| TCGA-AB-2899 | female | 76 | 22.4 | dead |
| TCGA-AB-2900 | male | 70 | 6.1 | dead |
| TCGA-AB-2901 | male | 27 | 2.0 | alive |
| TCGA-AB-2908 | male | 81 | 1.0 | dead |
| TCGA-AB-2911 | female | 51 | 39.5 | alive |
| TCGA-AB-2912 | male | 63 | 9.1 | dead |
| TCGA-AB-2913 | male | 61 | 40.5 | alive |
| TCGA-AB-2914 | female | 22 | 26.4 | alive |
| TCGA-AB-2916 | female | 48 | 29.4 | alive |
| TCGA-AB-2917 | female | 41 | 40.5 | alive |
| TCGA-AB-2919 | female | 54 | 2.0 | alive |
| TCGA-AB-2920 | male | 44 | 12.2 | dead |
| TCGA-AB-2924 | male | 59 | 3.0 | alive |
| TCGA-AB-2925 | male | 57 | 8.1 | dead |
| TCGA-AB-2927 | female | 88 | 3.0 | dead |
| TCGA-AB-2928 | female | 43 | 2.0 | dead |
| TCGA-AB-2929 | female | 71 | 4.1 | dead |
| TCGA-AB-2933 | male | 58 | 4.1 | dead |
| TCGA-AB-2934 | male | 65 | 0.9 | alive |
| TCGA-AB-2935 | male | 66 | 2.0 | dead |
| TCGA-AB-2936 | female | 61 | 2.0 | alive |
| TCGA-AB-2937 | female | 36 | 7.2 | dead |
| TCGA-AB-2939 | male | 72 | 15.2 | alive |
| TCGA-AB-2942 | female | 67 | 21.4 | alive |
| TCGA-AB-2948 | male | 67 | 19.3 | dead |
| TCGA-AB-2949 | male | 58 | 23.3 | alive |
| TCGA-AB-2950 | female | 34 | 10.2 | alive |
| TCGA-AB-2952 | female | 60 | 1.0 | dead |
| TCGA-AB-2955 | female | 56 | 16.3 | dead |
| TCGA-AB-2956 | male | 61 | 6.1 | dead |
| TCGA-AB-2959 | male | 71 | 16.3 | dead |
| TCGA-AB-2963 | male | 56 | 54.7 | dead |
| TCGA-AB-2965 | male | 60 | 11.2 | dead |
| TCGA-AB-2966 | female | 57 | 28.5 | dead |
| TCGA-AB-2970 | female | 34 | 10.2 | dead |
| TCGA-AB-2971 | female | 76 | 26.4 | dead |
| TCGA-AB-2973 | female | 68 | 20.3 | dead |
| TCGA-AB-2976 | male | 53 | 30.5 | dead |
| TCGA-AB-2977 | female | 71 | 1.0 | dead |
| TCGA-AB-2979 | female | 30 | 22.4 | alive |
| TCGA-AB-2980 | male | 50 | 23.3 | alive |
| TCGA-AB-2981 | female | 35 | 16.2 | alive |
| TCGA-AB-2982 | female | 29 | 5.0 | alive |
| TCGA-AB-2983 | male | 45 | 11.2 | dead |
| TCGA-AB-2984 | male | 38 | 38.6 | alive |
| TCGA-AB-2986 | female | 31 | 7.1 | dead |
| TCGA-AB-2987 | female | 75 | 6.1 | dead |
| TCGA-AB-2988 | female | 67 | 1.0 | dead |
| TCGA-AB-2990 | male | 51 | 15.2 | alive |
| TCGA-AB-2991 | female | 40 | 60.9 | alive |
| TCGA-AB-2992 | female | 32 | 56.9 | dead |
| TCGA-AB-2995 | male | 63 | 51.7 | alive |
| TCGA-AB-2996 | male | 74 | 52.7 | alive |
| TCGA-AB-2998 | female | 68 | 1.0 | dead |
| TCGA-AB-2999 | male | 62 | 57.8 | alive |
| TCGA-AB-3001 | female | 31 | 52.7 | alive |
| TCGA-AB-3002 | male | 68 | 47.7 | dead |
| TCGA-AB-3007 | male | 35 | 52.7 | alive |
| TCGA-AB-3008 | male | 22 | 27.4 | dead |
| TCGA-AB-3009 | male | 23 | 19.2 | dead |
| TCGA-AB-3011 | female | 21 | 62.8 | alive |
| TCGA-AB-3012 | male | 53 | 62.9 | alive |

**Table S2.** 98 survival-related immune genes (SIGs).

| **SIGs** | **p-value** |
| --- | --- |
| ADAM12 | 0.009055077 |
| ADAMTS3 | 0.019672714 |
| ADGRE5 | 0.030478533 |
| AHNAK | 0.018578223 |
| AIF1 | 0.01115535 |
| AKT3 | 0.029314184 |
| ANXA5 | 0.041670163 |
| APOL3 | 0.012945343 |
| ARHGAP10 | 0.003128417 |
| ARL1 | 0.019752628 |
| ASB2 | 0.005555865 |
| ATP6V1A | 0.040023492 |
| BST2 | 0.017630636 |
| C1orf162 | 0.024263467 |
| C1orf54 | 0.014255787 |
| CCR5 | 0.013876728 |
| CD109 | 0.012251905 |
| CD37 | 0.000151561 |
| CD4 | 0.013047345 |
| CD52 | 0.000148262 |
| CD79B | 0.047103955 |
| CFL1 | 0.028244461 |
| CLCN5 | 0.0000025 |
| CLIC2 | 0.012485077 |
| CLIC3 | 0.025449448 |
| CREB5 | 0.024746533 |
| CRYBB1 | 0.031592817 |
| CSF2RA | 0.040604671 |
| CTPS1 | 0.006880889 |
| CTSD | 0.016154468 |
| CXCR2 | 0.039440985 |
| CYP27A1 | 0.032368574 |
| CYTH1 | 0.010866987 |
| DAB2 | 0.030817755 |
| DAPK1 | 0.046970159 |
| DARS | 0.009274375 |
| DAXX | 0.000084 |
| DPYD | 0.013317327 |
| EMP3 | 0.007094884 |
| F12 | 0.03259722 |
| FABP5 | 0.004115137 |
| FAM49A | 0.021102459 |
| FERMT3 | 0.000482707 |
| GAB3 | 0.012790197 |
| HAL | 0.030493772 |
| HCP5 | 0.005447746 |
| HELLS | 0.030883597 |
| HNMT | 0.016554784 |
| HRH1 | 0.015225832 |
| HSPA6 | 0.030522513 |
| ICAM3 | 0.048885585 |
| IDH3A | 0.034302608 |
| IGFBP5 | 0.031599509 |
| IKZF1 | 0.03136533 |
| IL3RA | 0.030153684 |
| INPP5F | 0.003711848 |
| ITGAL | 0.002788804 |
| ITGAM | 0.036646269 |
| ITGB2 | 0.030734879 |
| ITGB4 | 0.020982102 |
| KIR2DL1 | 0.011560605 |
| KIR3DL2 | 0.048387167 |
| KLRG1 | 0.015460397 |
| LAMP2 | 0.013655645 |
| LGALS1 | 0.000181036 |
| LRRC42 | 0.020273562 |
| LSP1 | 0.000744407 |
| LST1 | 0.015009468 |
| MAK | 0.030356179 |
| MBP | 0.038905732 |
| MMD | 0.017479911 |
| MOCOS | 0.004815371 |
| NCAM1 | 0.009065868 |
| NUCB2 | 0.00788809 |
| OFD1 | 0.022159756 |
| PDCD1 | 0.021997507 |
| PIK3IP1 | 0.006985001 |
| PLA2G4A | 0.033592713 |
| PNPLA6 | 0.007064373 |
| PTGES2 | 0.035224203 |
| PTGS1 | 0.002302456 |
| RPS9 | 0.026383806 |
| RUBCN | 0.001673805 |
| S100A4 | 0.000139415 |
| SELE | 0.004005489 |
| SFXN3 | 0.001168979 |
| SIGLEC10 | 0.036557072 |
| SLA | 0.027371484 |
| SPCS3 | 0.033123296 |
| ST3GAL6 | 0.022176552 |
| ST8SIA4 | 0.045818496 |
| TACSTD2 | 0.025528048 |
| TLR9 | 0.03758144 |
| TNFAIP2 | 0.001039854 |
| TOGARAM2 | 0.030739985 |
| TOX4 | 0.034453285 |
| UPP1 | 0.046621377 |
| VCAM1 | 0.030889137 |

